# Reduction Midpoint Potential of a Paradigm Light-Oxygen-Voltage Receptor and its Modulation by Methionine Residues

**DOI:** 10.1101/2024.02.29.582800

**Authors:** Andrés García de Fuentes, Andreas Möglich

## Abstract

Light-dependent adaptations of organismal physiology, development, and behavior abound in nature and depend on sensory photoreceptors. As one class, light-oxygen-voltage (LOV) photoreceptors harness flavin-nucleotide chromophores to sense blue light. Photon absorption drives the LOV receptor to its signaling state, characterized by a metastable thioadduct between the flavin and a conserved cysteine residue. With this cysteine absent, LOV receptors instead undergo photoreduction to the flavin semiquinone which however can still elicit downstream physiological responses. Irrespective of the cysteine presence, the LOV photochemical response thus entails a formal reduction of the flavin. Against this backdrop, we here investigate the reduction midpoint potential *E*_0_ in the paradigm LOV2 domain from *Avena sativa* phototropin 1 (*As*LOV2), and how it can be deliberately varied. Replacements of residues at different sites near the flavin by methionine consistently increase *E*_0_ from its value of around –280 mV by up to 40 mV. Moreover, methionine introduction invariably impairs photoactivation efficiency and thus renders the resultant *As*LOV2 variants less light-sensitive. Although individual methionine substitutions also affect the stability of the signaling state and downstream allosteric responses, no clear-cut correlation with the redox properties emerges. With a reduction midpoint potential near –280 mV, *As*LOV2 and, by inference, other LOV receptors may be partially reduced inside cells which directly affects their light responsiveness. The targeted modification of the chromophore environment, as presently demonstrated, may mitigate this effect and enables the design of LOV receptors with stratified redox sensitivities.

## Introduction

Across diverse organisms, sensory photoreceptor proteins afford the sensation of light^1,2^. To this end, photoreceptors bind aromatic molecules as chromophores, thus enabling them to detect specific bands within the near-ultraviolet to near-infrared portion of the electromagnetic spectrum. Light absorption ushers in a photocycle, i.e., a series of (photo)chemical reactions, that culminate in population of the so-called signaling state which differs in its biological activity from the original, resting state prevailing in darkness. Light signals are thereby converted into biological responses with great precision in space and time, and in reversible manner. Beyond their pivotal roles in nature, photoreceptors have garnered additional prominence as the key ingredients in optogenetics^3^: Introduced to heterologous organisms by genetic means, photoreceptors allow the unprecedented light-dependent control and perturbation of cellular state and processes. Originating in the neurosciences, optogenetics has long transcended these beginnings and now widely applies to diverse use cases in molecular and cell biology^4,5^.

The above aspects are exemplified by light-oxygen-voltage (LOV) proteins, a class of photoreceptors binding flavins in their oxidized quinone state as chromophores^6–8^. First discovered in plants^6,7^, but also widespread in bacteria^9^ and fungi, LOV receptors traverse a distinct photocycle upon blue-light absorption^10^. An initially formed, electronically excited singlet state (S_1_) of the flavin chromophore rapidly transitions to a triplet state (T_1_) by intersystem crossing (Fig. 1a). The T_1_ state converts into the signaling state that is characterized by a covalent adduct between the C4a atom of the flavin isoalloxazine and the Sγ atom of a close-by, conserved cysteine residue^11^. This reaction likely proceeds via a radical-pair intermediate^12,13^ and amounts to a formal reduction of the flavin. Within the adduct state, the N5 atom is thus protonated and acts as a hydrogen-bond donor, rather than an acceptor as in the resting receptor state. As best characterized for the paradigm LOV2 domain of *Avena sativa* phototropin 1 (*As*LOV2)^14–16^ (Fig. 1b), said protonation induces hydrogen-bonding rearrangements of a proximal glutamine and downstream residues that jointly underlie the light-dependent regulation of receptor activity. The thioadduct spontaneously reverts to the resting state in the base-catalyzed^17^ dark-recovery reaction. Despite strong conservation in LOV receptors, the active-site cysteine is not essential for blue-light-dependent signaling^18^, nor is the glutamine^16^. In the absence of the cysteine, the absorption of blue light can prompt the reduction of the flavin to the semiquinone (SQ) radical state^19,20^. Given its high pKa value, the N5 atom is readily protonated, and the neutral SQ (NSQ) forms which is capable of downstream signaling^18^.

**Figure 1.**
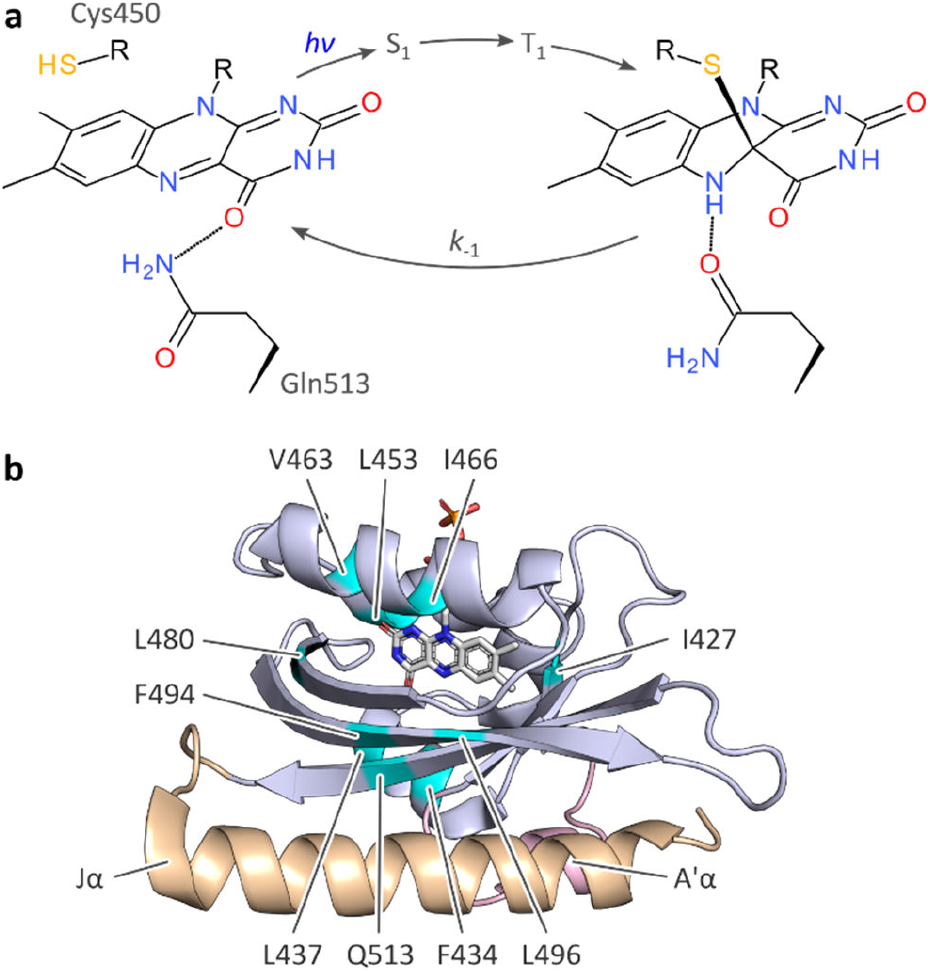
Photocycle and structure of the *A. sativa* phototropin 1 LOV2 (*As*LOV2) domain. **a**, Light-oxygen-voltage (LOV) receptors absorb blue light via flavin-nucleotide chromophores in their oxidized quinone state. Upon photoexcitation, the flavin undergoes rapid intersystem crossing from a singlet (S_1_) to a triplet state (T_1_), out of which the thioadduct signaling state forms. Owing to the covalent adduct between a conserved cysteine residue and the C4a atom of the isoalloxazine, the flavin N5 position is protonated which is key to downstream signal transduction^11,14,16^. **b**, Structure of *As*LOV2 in its dark-adapted state (pdb entry 7pgx^16^), with the LOV core domain and the A’α and Jα helices shown in blue, rose, and orange, respectively. Residues at which methionine residues were introduced are highlighted in cyan.

Considered as a whole, signal transduction in LOV receptors hinges on flavin reduction and protonation, both in the presence and absence of the cysteine. Hence, the redox environment may affect various aspects of the LOV signaling trajectory. For instance, chemical reduction of the flavin quinone state, usually a two-electron reduction^21^, to the hydroquinone entails N5 protonation and may hence trigger downstream signal transduction. In fact, it has been proposed that certain LOV receptors may double as cellular redox sensors^22,23^. Previous studies put the midpoint potentials of LOV receptors in the range of –250 mV to –300 mV^17,21–23^. There is only scarce data on how the redox properties of LOV receptors affect signal transduction, but one report found little effect of the reduction potential on LOV dark recovery, whereas other parameters await in-depth probing^21^. The targeted modulation of the reduction midpoint potential in LOV receptors is similarly unexplored. The removal of the active-site cysteine led to a 10 mV higher, i.e., less negative, flavin reduction midpoint potential in a phototropin LOV domain^21^. Within the LovK histidine kinase from *Caulobacter crescentus*, serial truncations of the α-helical linker C-terminal of the LOV sensor module prompted sizeable shifts in the reduction midpoint potential^22^, but a later study failed to reproduce these findings^24^. In the short-LOV receptor Vivid from *Neurospora crassa* (*Nc*VVD), replacement by isoleucine of two methionines near the flavin chromophore incurred partial reduction to the NSQ state during heterologous expression in *Escherichia coli*^18,25^. This observation may signify a higher flavin reduction midpoint potential caused by the removal of two electron-rich sulfur atoms^18,25^.

Piqued by these reports, we here investigate the redox properties of the paradigm *As*LOV2 module, and to which extent it can be modulated. We find that the introduction of methionines at several sites around the chromophore indeed changes the flavin reduction midpoint potential by up to 40 mV. Contrary to the proposal for *Nc*VVD^18,25^, the presence of methionines however incurs an increase in midpoint potential rather than a decrease. Irrespective of where within *As*LOV2 the methionines are placed, light-induced thioadduct formation becomes substantially less efficient. By contrast, no systematic effect on the dark-recovery reaction manifests, nor on the flavin fluorescence. Although downstream signaling is impaired for certain methionine insertion sites, it remains intact for other positions. This opens the prospect of devising receptors that integrate light and redox cues and process them into biological responses.

## Results and discussion

### Design of *As*LOV2 methionine variants

To probe the effect of methionine residues on reduction midpoint potential and other aspects of LOV receptors, we opted for the *As*LOV2 module as a model system, given its well-developed photochemical^26,27^, structural^14–16^, and mechanistic characterization, and its pervasive use as a versatile building block in optogenetics^4,28^. Based on previous reports^18,29–31^ and structural inspection, we selected 13 inward-facing, predominantly hydrophobic residues around the flavin mononucleotide (FMN) cofactor for replacement by methionine, including isoleucine I466 and leucine L496 that structurally correspond to the methionine sites M135 and M165 investigated in *Nc*VVD^25^ (Fig. 1b). Upon heterologous expression in *E. coli*, 10 *As*LOV2 variants with a methionine at the residue positions I427, F434, L437, L453, V463, I466, L480, F494, L496, and Q513 could be produced in soluble form and sufficient quantity. By contrast, methionine introduction at the positions V416 and G511 failed as no soluble holoprotein could be isolated. We additionally generated one variant that combines the I466M and L496M exchanges.

### Methionine substitutions modulate LOV photocycle kinetics

As analyzed by UV-vis absorbance spectroscopy, wild-type *As*LOV2 and the 11 methionine variants that could be produced in soluble form (Table 1) incorporated their flavin chromophores in the oxidized quinone state (Fig. 2a, Suppl. Fig. 1). Likewise, all the tested *As*LOV2 variants retained the ability to form the cysteinyl:flavin thioadduct state under blue light, as indicated by the disappearance of the three-pronged absorbance peak around 450 nm of the dark-adapted state and the emergence of a broader absorbance band peaking at 390 nm and characteristic of the light-adapted signaling state. In the variants I427M, F434M, and L453M, the light-adapted state was not populated to full extent, and a minor portion of the dark-adapted species persisted even after prolonged illumination. We ascribe these observations to a moderately sped up dark recovery compared to wild type (see below). These aspects aside, the shape and location of the absorbance spectra prior to and after light exposure were closely similar across all variants, with Q513M being the only exception (Fig. 2d). As reported earlier^16,32,33^, replacement of this glutamine, in the past often by leucine, induces a hypsochromic spectral shift of around 6 nm in both the dark-adapted and light-adapted states which can be tied to a loss of hydrogen bonding to the flavin O4 atom.

**Table 1.** Properties of *As*LOV2 methionine variants.

| AsLOV2 variant <sup>a</sup> | dark recovery <sup>b</sup> | | activation <sup>c</sup> | circular dichroism <sup>b</sup><br>[$\theta$]<sub>MRW</sub><sup>208</sup> × 10<sup>-3</sup><br>(deg cm<sup>2</sup> dmol<sup>-1</sup>) | | fluorescence <sup>b</sup> | midpoint potential <sup>b</sup> |
| --- | --- | --- | --- | --- | --- | --- | --- |
| | *k*<sub>-1</sub> (s<sup>-1</sup>) | *E*<sub>a</sub> (kJ mol<sup>-1</sup>) | *k*<sub>1</sub> (s<sup>-1</sup>) | dark | light | $\Phi_f$ | *E*<sub>0</sub> (mV) |
| wild type | 0.013 | 61.0 | 0.048 | -12.0 | -9.0 | 0.09 | -278 |
| I427M | 0.050 | 70.5 | 0.018 | -13.5 | -11.9 | 0.10 | -260 |
| F434M | 0.024 | 43.8 | 0.021 | -12.7 | -10.2 | 0.08 | -258 |
| L437M | 0.011 | 59.5 | 0.027 | -13.4 | -9.4 | 0.08 | -263 |
| L453M | 0.047 | 61.4 | 0.032 | -12.4 | -10.2 | 0.09 | -268 |
| V463M | 0.014 | 84.3 | 0.016 | -9.9 | -9.1 | 0.09 | -236 |
| I466M | 0.012 | 64.7 | 0.039 | -13.5 | -10.5 | 0.07 | -257 |
| L480M | 0.020 | 32.9 | 0.026 | -10.7 | -10.3 | 0.04 | -270 |
| F494M | 0.003 | 70.4 | 0.013 | -12.1 | -10.5 | 0.04 | -268 |
| L496M | 0.018 | 63.8 | 0.028 | -10.8 | -7.1 | 0.08 | -279 |
| Q513M | 0.002 | 62.9 | 0.005 | -14.1 | -13.1 | 0.04 | -252 |
| I466M:L496M | 0.019 | 65.0 | 0.019 | -12.8 | -10.3 | 0.07 | -263 |

<sup>a</sup>: All AsLOV2 variants comprise residues 404–546.
<sup>b</sup>: Determined at 22 °C.
<sup>c</sup>: Measured at 22 °C for a set illumination geometry and a blue-light intensity of 3 mW cm<sup>-2</sup>.

**Figure 2.**
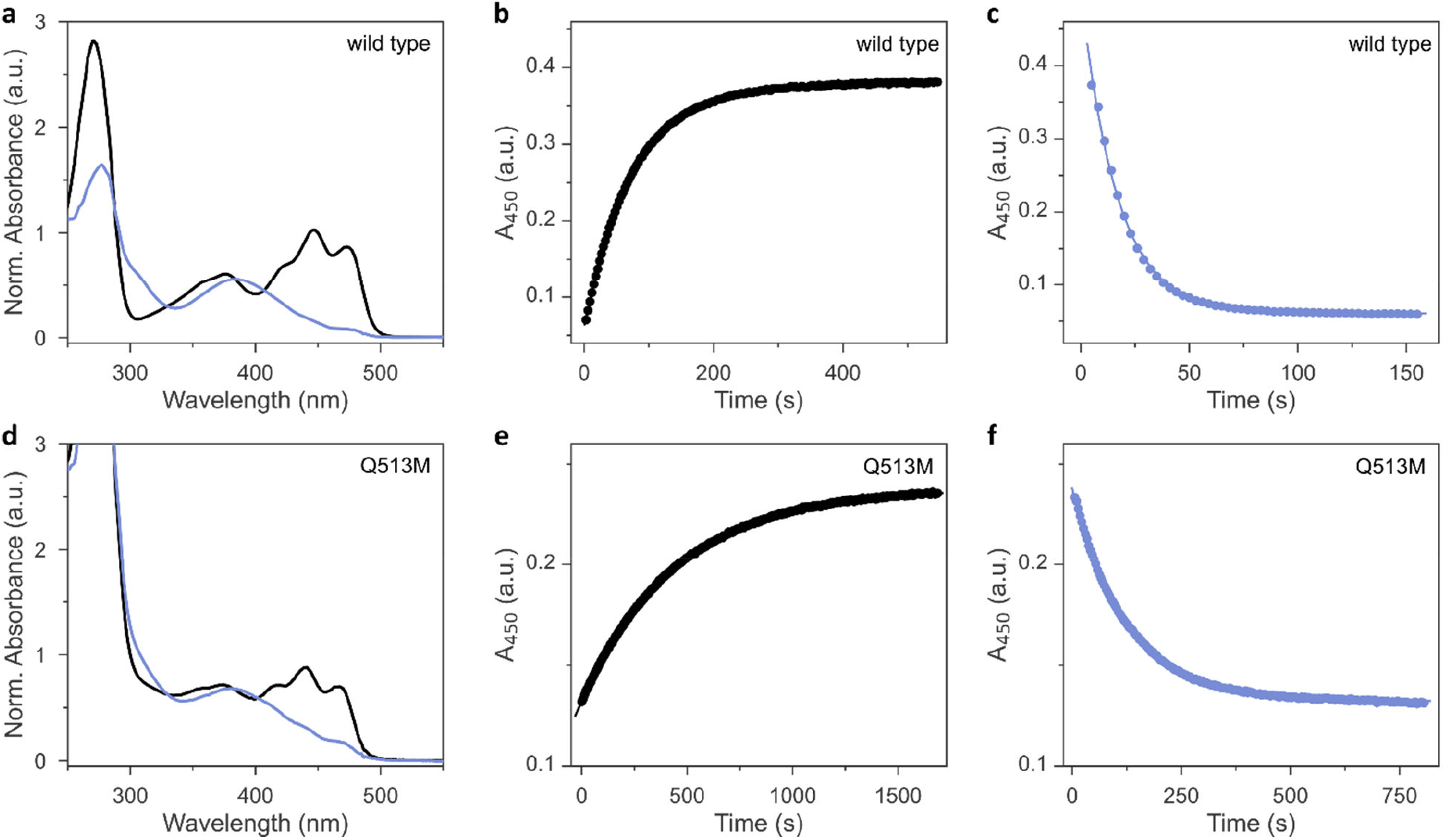
Photoactivation and dark recovery of *As*LOV2 variants. **a**, UV-vis absorbance spectra of *As*LOV2 wild type in its dark-adapted (black lines) and light-adapted states (blue). **b**, Dark recovery after saturating illumination with blue light, monitored at 450 nm. **c**, Photoactivation kinetics were recorded by exposing dark-adapted *As*LOV2 to 3 mW cm^-2^ blue light, while tracking the transition to the light-adapted state at 450 nm. **d-f**, As panels a-c but for *As*LOV2 Q513M.

We next assessed the dark-recovery reaction, i.e., the return of the LOV protein to the dark-adapted resting state once illumination ceases (Fig. 2b, Suppl. Fig. 2). At 22°C, wild-type *As*LOV2 reverted to its resting state in a single-exponential process with a rate constant *k*_-1_ of 0.013 s^-1^. Generally, the recovery strongly depends on the packing of (mostly) hydrophobic residues within the LOV core domain and solvent accessibility to the flavin. Many residue exchanges have been identified that modulate the velocity of the recovery kinetics over several orders of magnitude^34^. All 12 *As*LOV2 variants tested at present exhibited complete recovery to the fully dark-adapted state. The variants I427M and L453M sped up the dark reversion by more than 3-fold, in line with earlier analyses showing these residue positions to affect the recovery reaction^34,35^. Contrarily, the dark recovery slowed down by more than 4-fold in the F494M and Q513M variants (Fig. 2e, Suppl. Fig. 2), again consistent with earlier reports which implicated these residue positions in affecting dark recovery^34^. The strongest deceleration was evidenced for Q513M and can be attributed to the removal of the amide sidechain of the glutamine, thus hindering N5 deprotonation and rupture of the thioadduct during dark reversion to the resting state^16,32–34^. Based on recovery experiments conducted at temperatures from 22°C to 38°C, we calculated Arrhenius activation energies for the dark recovery which amounted to 61 kJ mol^-1^ for wild-type *As*LOV2 (Table 1). Most *As*LOV2 variants had similar activation energies but those of F434M and L480M were significantly lower at 44 kJ mol^-1^ and 33 kJ mol^-1^, respectively. Although a molecular rationalization is elusive, the lower activation energies hint at mechanistic differences in the dark-recovery process in these two variants.

### Methionine residues desensitize LOV adduct formation

Next, we reasoned that placing a sulfur-containing methionine sidechain close-by the flavin chromophore may well impact on the photochemical events playing out upon photoexcitation. To check that possibility, we investigated the photoactivation kinetics with which the dark-adapted quinone state of the *As*LOV2 variants converts to the thioadduct signaling state under continuous illumination (Fig. 2c). As noted above, en route to the adduct state, the LOV receptor traverses the S_1_ and T_1_ electronically excited states and likely a transient radical-pair intermediate^10^. The overall quantum yield *Φ*_390_ for this reaction series leading to thioadduct formation amounts to between 0.3 and 0.5 for phototropin LOV2 domains in general^36^, and to around 0.44 for *As*LOV2 in particular^37^. Using absorbance spectroscopy, we monitored how fast relative to the wild type the individual *As*LOV2 variants acquire their signaling state when exposed to a blue light-emitting diode (LED) (Table 1, Suppl. Fig. 3). For this comparison, it is imperative that the illumination conditions be consistent which we achieved by careful light power calibration and by using a 3D-printed adapter for precise mounting of the LED. With a set illumination geometry and light power (450 nm, 3 mW cm^-2^), the wild-type protein assumed its signaling state in a single-exponential reaction with a rate constant of 0.062 s^-1^ (Fig. 2c). By accounting for the recovery simultaneously taking place during illumination, we calculated a unimolecular rate constant of 0.048 s^-1^ for photoactivation under the given experimental conditions. Intriguingly, all the *As*LOV2 variants reached their signaling state more slowly under the same illumination settings (Suppl. Fig. 3). As their absorbance spectra did not substantially differ from that of the wild-type protein (see Fig. 2 and Suppl. Fig. 1), we interpret these data as a reduction of *Φ*_390_ in the *As*LOV2 methionine variants. Put another way, the corresponding variants are less light-sensitive than the wild-type receptor. The effect was most pronounced for the Q513M substitution which gave rise to almost tenfold slower photoactivation kinetics (Fig. 2f).

**Figure 3.**
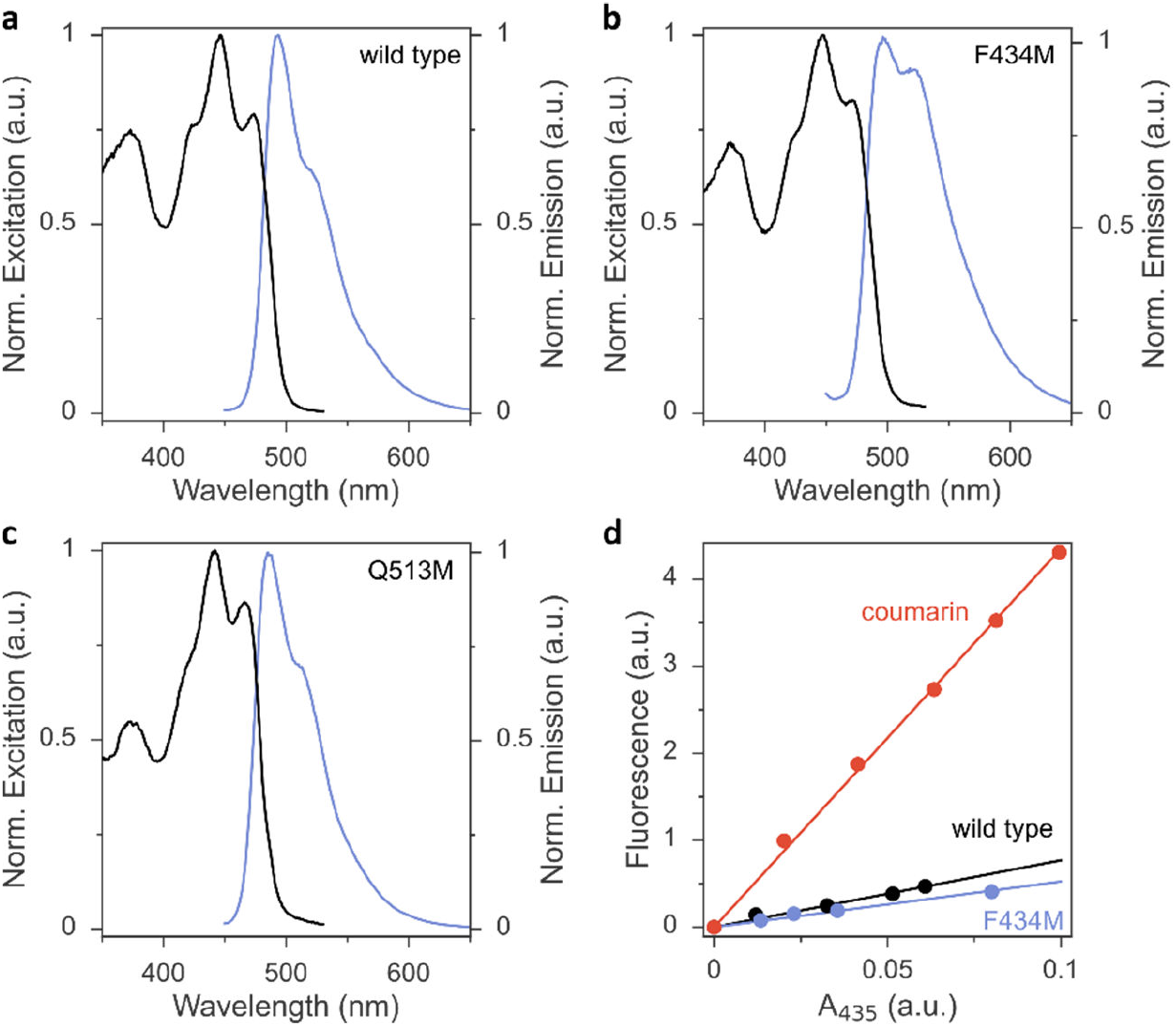
Fluorescence excitation (black lines) and emission spectra (blue) of *As*LOV2 variants. **a**, Wild-type *As*LOV2. **b**, *As*LOV2 F434M. **c**, *As*LOV2 Q513M. **d**, Determination of fluorescence quantum yield *Φ*_F_ in *As*LOV2 wild type (black symbols) and F434M (blue) by comparison to coumarin 153 (red) with known *Φ*_F_ of 0.54. The abscissa and ordinate denote the absorbance at the excitation wavelength 435 nm and the integrated fluorescence, respectively.

Noted above, the modulation of LOV recovery kinetics by residue replacements is well developed^34^. A set of proven residue exchanges is at disposal for reliably altering these kinetics across various LOV receptors. Besides mechanistic interest, such residue substitutions grant immediate benefit for applications in optogenetics and biotechnology as they allow adjusting the effective light sensitivity of a photoreceptor at photostationary state under continuous illumination^38,39^. The faster the reversion to the resting state, the more light is needed to populate the signaling state, and the less light-sensitive a given photoreceptor circuit effectively becomes. By contrast, much less is known about how the absolute light sensitivity, governed by *Φ*_390_, can be varied at will. This is all the more surprising as substantial differences in absolute light sensitivity among phototropin LOV1 and LOV2 modules have been long documented^37^, if not mechanistically rationalized. Our work now identifies a means of deliberately lowering *Φ*_390_ and of thus partially desensitizing the LOV receptor to blue light. By contrast, we expect that raising *Φ*_390_ will prove exceedingly challenging as, at least in the case of LOV2 domains, the physically possible limit may have been *de facto* approached in the course of evolution already.

To glean additional insight into the effect of the methionine substitutions on *Φ*_390_, we assessed the flavin fluorescence in the *As*LOV2 proteins (Fig. 3, Suppl. Fig. 4). Most variants had closely similar emission spectra with a peak around 495 nm and a shoulder at around 520 nm. As an exception, the Q513M variant exhibited hypsochromic shifts of both its excitation and emission spectra, in line with the corresponding shift observed in the absorbance spectrum (Fig. 3c). Further, the F434M exchange led to an intensity increase of the emission around 520 nm (Fig. 3b), reminiscent of earlier reports that implicated this residue position in determining fluorescence properties.^40^ The fluorescence quantum yield *Φ*_F_ was 0.09 in wild-type *As*LOV2, consistent with earlier measurements^41^ (Table 1, Fig. 3d). Methionine introduction had minor effects on *Φ*_F_ of most variants. This hints at similar lifetimes of the excited S_1_ state which in turn suggests that the reduced *Φ*_390_ (see above) mainly originates from impaired thioadduct formation out of the T_1_ state. As an exception, the L480M, F494M, and Q513M species showed substantially reduced *Φ*_F_ of around 0.04, indicating that in these variants the lower *Φ*_390_ owes at least in part to a shortened S_1_ lifetime.

**Figure 4.**
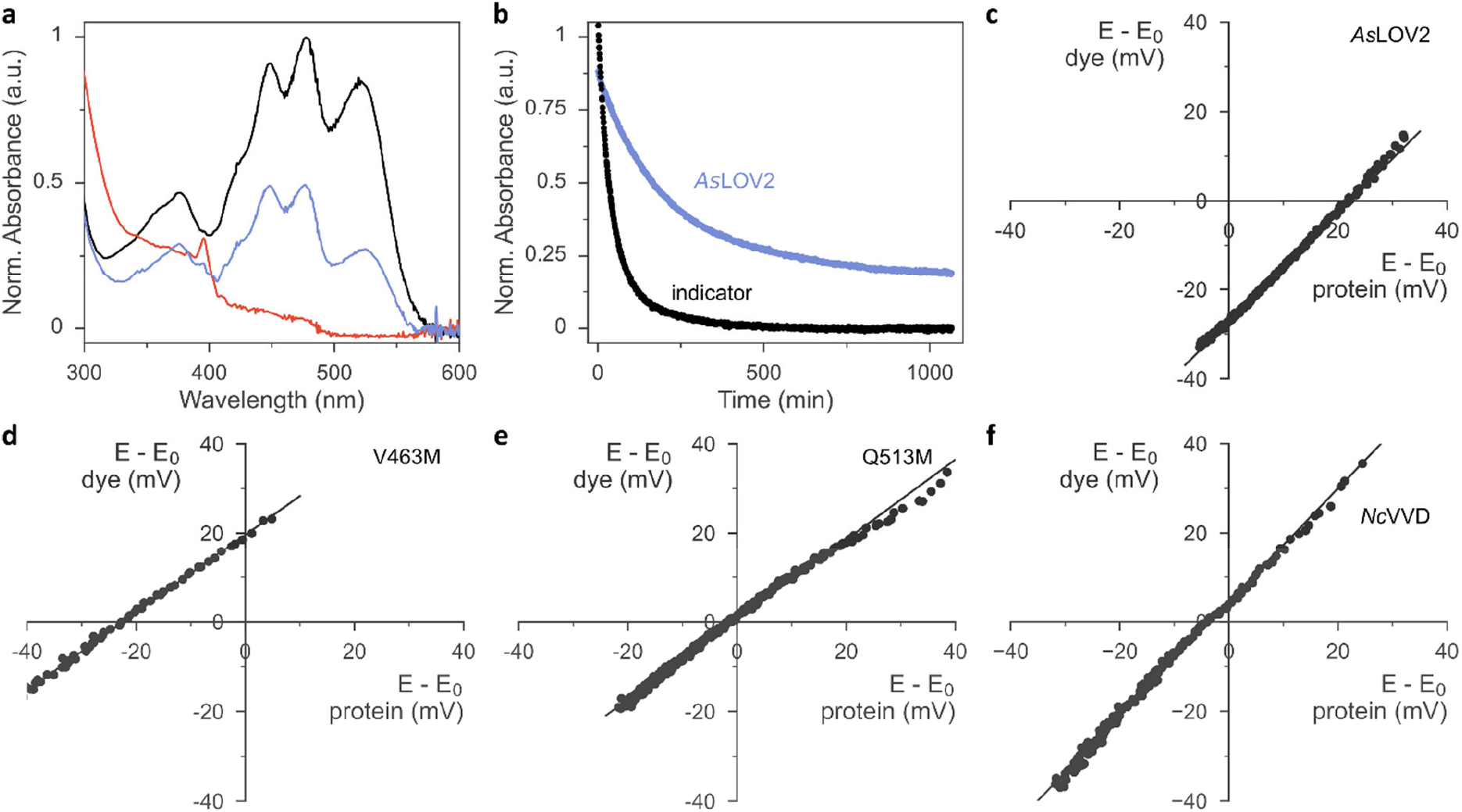
Determination of flavin reduction midpoint potential in LOV domains. **a**, UV-vis absorbance spectra of *As*LOV2 wild-type in the presence of the redox indicator phenosafranine and dithiothreitol (DTT). The spectrum shown in black was recorded immediately upon DTT addition, and the spectra shown in blue and red reflect timepoints 200 and 800 minutes afterwards, respectively. **b**, The absorbance reading at 520 nm reports on the reduction of the indicator over time (grey symbols). The absorbance signal at 450 nm (blue) was corrected for the contribution of the redox indicator and reports on the reduction of the *As*LOV2 protein. **c**, From the data in panel b, the fractions of oxidized and reduced indicator and *As*LOV2 are calculated for given timepoints. The difference in reduction midpoint potentials between indicator (*y* axis) and *As*LOV2 (*x* axis) is determined according to the Nernst equation. **d**, As panel c but for *As*LOV2 V463M. **e**, As panel c but for *As*LOV2 Q513M. **f**, As panel c but for *Nc*VVD.

H2>Methionine introduction increases the *As*LOV2 flavin reduction midpoint potential

Having ascertained the preservation of thioadduct formation, we next evaluated the reduction midpoint potentials *E*_0_ in *As*LOV2 wild type and its methionine variants. This potential can be measured by varying the reduction potential *E* in solution while tracking the fractions of oxidized and reduced analyte. For flavins, these fractions can be determined by spectroscopy as the oxidized quinone, the partially reduced semiquinone (SQ), and the fully reduced hydroquinone (HQ) states exhibit distinct absorbance spectra^21,24,42^. Earlier reports revealed that the chemical reduction of LOV proteins is usually a two-electron process and yields the HQ^21,22,24^, whereas photoreduction involves a one-electron transfer and produces singly reduced SQ radical states^18–20,43^. For the calculation of *E*_0_ according to the Nernst equation, the reduction potential *E* is inferred by monitoring the absorbance of a redox indicator dye of known midpoint potential, e.g., phenosafranine with *E*_r0_ = –252 mV. This value and all subsequent reduction midpoint potentials are relative to the normal hydrogen electrode. A key prerequisite for this analysis is that the system be in redox equilibrium. To this end, methyl viologen is included as a mediator substance that ferries electrons and facilitates equilibration between the analyte, the indicator dye, and the bulk solvent. A convenient way of recording a complete redox isotherm in one go is provided by gradual variation of the reduction potential *E* via an enzymatic reaction, most often the xanthine/xanthine oxidase (XO) system. Hydrogen peroxide, formed by XO, and molecular oxygen are commonly mopped up by added catalase and glucose oxidase.

Using this setup, we derived a value of –271 mV for the reduction midpoint potential of *As*LOV2 wild type (Suppl. Fig. 5). When implementing the XO approach, we however found it demanding to titrate the enzyme amount appropriately and reproducibly. Using too much enzyme resulted in overly fast reaction time courses that violated the above requirement of steady-state redox equilibration. By contrast, with too little enzyme, the reduction of analyte or dye remained incomplete^24^. Batch-to-batch variability of XO further compounded the issue and thus complicated the experiments. We overcame these challenges by doing away with XO-mediated enzymatic reduction and moving to chemical reduction with dithiothreitol (DTT)^44^. Under the exclusion of oxygen, DTT addition prompted the protracted yet complete reduction of both the flavin analyte and indicator dye within several hours. Control experiments with riboflavin as the analyte yielded a reduction midpoint potential *E*_0_ = –224 mV, which was highly reproducible across several experiments and in good agreement with the literature value of –220 mV at a pH of 7.0^45,46^.

**Figure 5.**
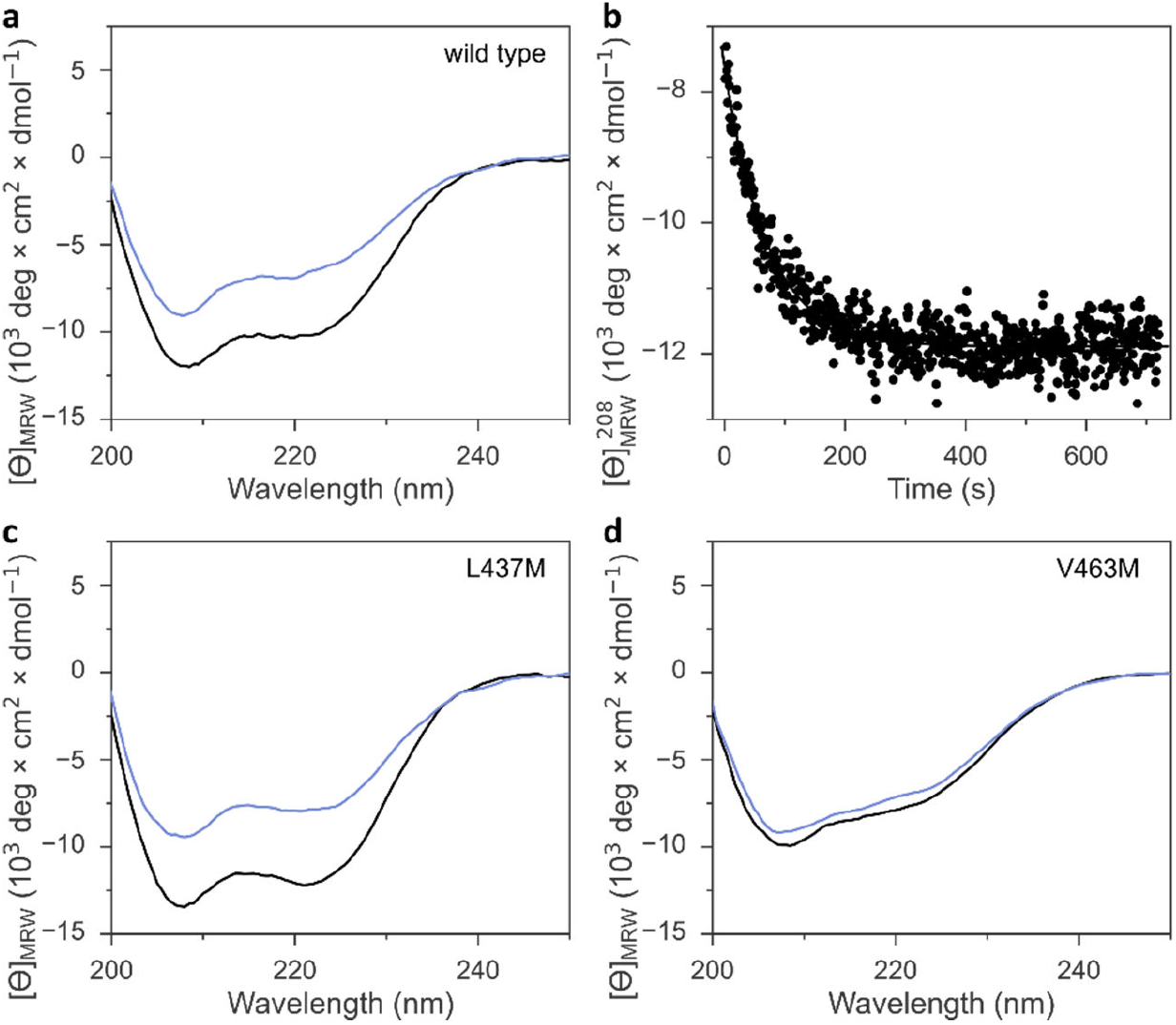
Signal transduction in *As*LOV2 variants. **a**, Far-UV circular dichroism (CD) spectra of *As*LOV2 in its dark-adapted (black line) and light-adapted states (blue). **b**, The recovery of *As*LOV2 to its dark-adapted state after blue-light exposure was monitored by tracking over time the CD signal at (208 ± 5) nm. **c**, As panel a but for *As*LOV2 L437M. **d**, As panel a but for *As*LOV2 V463M.

We next progressed to *As*LOV2 wild-type and observed complete reduction of the flavin cofactor to the hydroquinone state within several hours of DTT addition (Fig. 4a). By correlating these reduction kinetics with those of the indicator dye (Fig. 4b-c), we calculated the reduction midpoint potential of *As*LOV2 as –278 mV. We entertained the hypothesis that, at least in principle, the observed *As*LOV2 reduction kinetics may reflect slow protein unfolding and subsequent flavin reduction outside of the LOV protein rather than inside it. Several arguments compellingly argue against this scenario. First, the *E*_0_ value we determined for *As*LOV2 is in reasonable agreement with that determined by the XO method and well within the range of values previously reported for LOV receptors^21,22,24^. Second, the correlation of the DTT-driven reduction kinetics of analyte and indicator according to the Nernst equation yielded linear correlations with a slope near unity for *As*LOV2 and all its variants (Fig. 4c-e, Suppl. Fig. 6). Third, throughout the entire reaction time course, the flavin portion remaining in the oxidized state maintained the fine structure of its absorbance signal near 450 nm (see, e.g., Fig. 4a), indicative of binding by the LOV receptor. Fourth, in control experiments in which oxygen was admitted during the reaction, no reduction occurred. The *As*LOV2 domain remained folded throughout and the flavin chromophore bound, as again evidenced by the fine structure of the absorbance spectrum. Taken together, we conclude that the DTT-driven reduction kinetics genuinely report on the redox properties of *As*LOV2.

**Figure 6.**
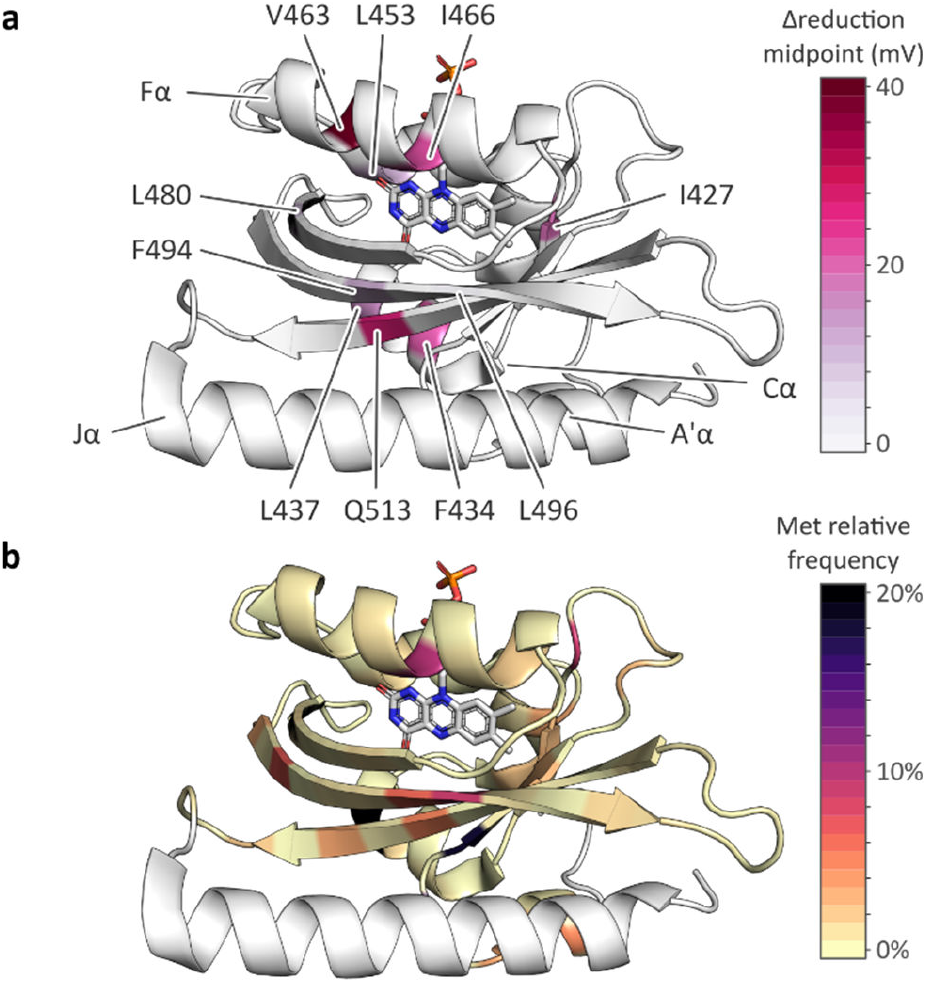
Effect of methionine introduction on *As*LOV2 reduction midpoint potential and methionine distribution within the light-oxygen-voltage (LOV) receptor family. **a**, Change in flavin reduction midpoint potential upon introduction of methionine residues at the indicated positions within *As*LOV2. **b**, Distribution of methionine residues within the light-oxygen-voltage (LOV) receptor family mapped onto the structure of *As*LOV2^16^. The color code denotes the relative frequencies of methionines at each residue position within a large-scale multiple sequence alignment of LOV receptors.

We next extended our studies to the *As*LOV2 methionine variants and consistently found *E*_0_ increases, except for the L496M substitution (Table 1, Fig. 4d-e, Suppl. Fig. 6). On average, methionine introduction incurred a 17 mV higher reduction midpoint potential, and this effect was most pronounced for the V463M variant with an *E*_0_ value of –236 mV (Fig. 4d). We next correlated the observed changes in reduction midpoint potential with the photochemical characteristics determined for the *As*LOV2 variants above. As noted above, methionines compromised photoactivation to the thioadduct state, almost irrespective of where within *As*LOV2 they were inserted. Beyond that, no clear-cut correlation between reduction midpoint potential and other parameters emerges.

The higher *E*_0_ values evidenced in the *As*LOV2 methionine variants run counter to the initial proposal formulated for *Nc*VVD that the removal of methionines, rather than their introduction, would increase the flavin reduction midpoint potential^18,25^. To resolve this conundrum, we prepared and analyzed several *Nc*VVD proteins. Apart from wild-type *Nc*VVD (residues 37–186), we investigated variants with the residues M135 and M136 individually or jointly replaced by isoleucine. The *Nc*VVD variant bearing both the substitutions M135I and M165I is termed VVD-II as previously^18,25^. Lastly, we also prepared the VVD-III variant that derives from VVD-II and replaces the active-site cysteine 108 by alanine. In wild-type *Nc*VVD, the reduction midpoint potential *E*_0_ was –240 mV, i.e., almost 40 mV higher than in *As*LOV2 (Table 2, Fig. 4f). The molecular underpinnings for this large difference remain elusive at present. Surprisingly, neither the single nor the double methionine removal led to significant changes in *E*_0_ of *Nc*VVD (Table 2, Suppl. Fig. 7). By contrast, a 20 mV decrease of midpoint potential resulted upon cysteine replacement in VVD-III. It is worth calling to mind that for a given reduction potential *E*, a lower reduction midpoint potential *E*_0_ favors the oxidized quinone over the reduced hydroquinone state of the flavin.

**Table 2.** Reduction midpoint potentials of *Nc*VVD variants.

| NcVVD variant <sup>a</sup> | midpoint potential <sup>b</sup><br>*E*<sub>0</sub> (mV) |
| --- | --- |
| wild type | -241 |
| M135I | -242 |
| M165I | -247 |
| M135I:M165I<br>(≡ VVD-II) | -243 |
| C108A:M135I:M165I<br>(≡ VVD-III) | -264 |

<sup>a</sup>: All NcVVD variants comprise residues 37–186.
<sup>b</sup>: Determined at 22 °C.

Our data accordingly show that the earlier assertion^18,25^ that methionine removal would prompt an increase of *E*_0_ does not hold up. This immediately raises the question of how then to account for the larger NSQ population seen in the VVD-II and especially the VVD-III variants upon their heterologous production in *E. coli*^18,25^. Although we can but speculate at present, we advance two conceivable scenarios. First, reductants within *E. coli* may serve as electron donors and thereby facilitate the one-electron reduction of the flavin quinone to the (neutral) semiquinone. The differences in NSQ population across the *Nc*VVD variant series may have less to do with the flavin reduction midpoint potential, unchanged in VVD-II and moderately affected in VVD-III, but more with the accessibility of the hypothetical reductant to the chromophore-binding pocket. Second, we note that earlier reports consistently attributed SQ formation to photoreduction^19,20^, whereas the chemical reduction invariably gave rise to the hydroquinone, at least for the experimental conditions investigated. Potentially, the appearance of the NSQ state during the *Nc*VVD protein expression may hence be down to photoreduction as opposed to chemical reduction. This hypothesis finds support in a recent report that revealed a strong impact of methionine residues on photoreduction to the anionic and neutral SQ states in the very same *Nc*VVD receptor^43^.

### Impact of methionine substitutions on signal transduction

Irrespective of the methionine insertion site, the present *As*LOV2 variants retained light-induced thioadduct formation. Past analyses however revealed that residue exchanges near the flavin chromophore may well induce the desired effect, e.g., modulation of photocycle kinetics, but at the same time inadvertently impair signal transduction^47^. We hence probed the *As*LOV2 methionine variants by circular dichroism (CD) spectroscopy for their capability of transducing downstream signals. Signal transduction in *As*LOV2 revolves around the reversible unfolding of N-terminal and C-terminal helices, denoted A’α and Jα, under blue light which can be tracked by CD measurements^14^. In darkness, the wild-type protein adopts a CD spectrum with two minima around 208 and 222 nm, consistent with its mixed αβ protein fold. Light prompted an approximately 40% reduction in the molar ellipticity per residue, [*θ*]_MRW_, at 208 nm (Table 1, Fig. 5a). Returned to darkness (Fig. 5b), the CD spectrum fully recovered to the original state with kinetics closely matching those for the UV-vis absorbance signal at 450 nm that monitors the flavin thioadduct rupture (see Fig. 2b).

Most *As*LOV2 methionine variants showed CD spectra in darkness overall similar to that of the wild type (Table 1, Fig. 5c-d, Suppl. Fig. 8). However, *As*LOV2 V463M and L480M had significantly lower [*θ*]_MRW_, similar to that of the wild-type protein in the light-adapted state, which hints at constitutive unfolding of the terminal A’α and Jα helices in these variants, regardless of illumination (Fig. 5d). Most methionine variants retained light-induced, reversible CD changes, albeit often at reduced amplitude, e.g., in the variants I427M, F434M, L453M, and I466M, F494M, Q513M, and the doubly substituted I466M:I496M (Suppl. Fig. 8). The retention of light-induced CD changes in *As*LOV2 Q513M, though attenuated compared to wild type, is consistent with our recent finding that signaling responses are largely preserved in the YF1 and PAL LOV receptors upon replacement of their corresponding glutamines by leucine^16^. Without this glutamine, water molecules likely mediate light-dependent signal transduction instead^16^. Only very small light-induced differences in the CD spectrum emerged for the V463M and L480M variants. By contrast, the *As*LOV2 L437M and L496M variants maintained the light-induced CD response observed in the wild-type protein to full extent.

## Conclusions

### Methionines in LOV receptors

Our data reveal that, depending on position, methionine residues affect various traits of *As*LOV2. In almost all cases, the presence of methionines increased the flavin reduction midpoint potential by up to 40 mV (Table 1, Fig. 6a). The most pronounced changes occurred for methionines placed within the Cα (F434M and L437M) and Fα (V463M and I466M) helices, and when substituting the conserved glutamine Q513 within the Iβ strand by methionine. By contrast, methionine introduction at other sites within the *As*LOV2 domain prompted smaller effects. Beyond their effect on flavin reduction midpoint potential, the methionine residues generally lowered the efficiency of blue-light-induced thioadduct formation and downstream signaling responses. Judiciously placed methionine residues thus provide a means of deliberately desensitizing LOV receptors, which can for instance be capitalized on for applications in optogenetics^39,48^. By contrast, we detected no systematic impact on other parameters, e.g., photoactivation, fluorescence, and signal transduction, although individual *As*LOV2 variants did exhibit sizeable differences in these quantities compared to the wild-type domain. This observation indicates that the reduction midpoint potential in LOV domains may be carefully varied while keeping other traits intact.

*As*LOV2 methionine variants with preserved signal transduction are prime candidates for the construction of LOV-based optogenetic switches with altered photochemical and redox properties. This is arguably most interesting for the *As*LOV2 variants I427M, L437M, and I466M given their diverging photoactivation kinetics, dark recovery, and reduction midpoint potential compared to wild type (Table 1). By contrast, the *As*LOV2 methionine variants with impaired light-induced signal transduction are less suited for exerting optogenetic control, but they may come to bear in LOV receptors with their cysteines removed which exhibit elevated fluorescence and propensity of generating reactive oxygen species (ROS). Provided that the methionine effects translate to the cysteine-devoid context, fluorescence reporters and ROS generators with stratified redox sensitivity may be now devised.

Given the often-substantial effect of methionines in *As*LOV2, we analyzed their distribution across the LOV receptor family (Fig. 6b). While methionines occur with elevated frequency of up to ∼ 20% at several positions within the LOV core domain, strikingly most of these sites face outward and thus point away from the flavin chromophore. Notable exceptions are the residue positions corresponding to L437, I466, and L496 in *As*LOV2, probed at present, that face towards the flavin chromophore. The relative paucity of methionine residues in the interior of the LOV domain could have different origins. On the one hand, it may be structurally challenging to accommodate the methionine sidechain inside the tightly packed hydrophobic domain core. On the other hand, methionines in the LOV interior may have been selected against given their presently documented effects on light detection and signal transduction.

### LOV has potential

A crucial, yet often disregarded aspect for applications of flavin-based receptors, e.g., LOV but also cryptochrome^49^ and BLUF receptors^50^, is how their reduction midpoint potentials stack up against the redox environment in the cellular context of interest. Based on evaluating the fraction of reduced vs. oxidized glutathione, the *E. coli* cytosol was found reducing with reduction potentials in the range of –260 mV to –280 mV^51^. A wider range between –300 mV^52^ and –230 mV^51^ has been reported for mammalian cells, with substantially higher values within the secretory pathway, under oxidative stress, and during apoptosis. Our and previous studies^21–24^ put the flavin reduction midpoint potentials of LOV receptors in a similar range (Table 1), which raises the possibility that LOV receptors may not fully assume their oxidized quinone state, that is commonly studied *in vitro*, but partially dwell in their reduced hydroquinone state inside cells. If so, LOV receptors may in principle moonlight as redox receptors as previously suggested^22–24^. The reduction to the hydroquinone state renders LOV and related receptors incapable of absorbing and reacting to blue light. Depending on the speed and extent of redox equilibration with the cellular environment, the light responsiveness of LOV-based circuits may be severely curtailed or lost altogether. Moreover, the response of a given circuit to blue light may vary depending on the subcellular compartment and whether the cell is experiencing oxidative stress. These considerations equally pertain to the use of derivative LOV modules as fluorescent reporters and ROS generators. We expect these aspects to be particularly relevant for LOV variants with a comparatively high reduction midpoint potential, e.g., *Nc*VVD.

In closing, we caution that cells rely on compartmentalized redox environments and multiple redox couples^52^. Cells generally operate far from chemical equilibrium and maintain distinct ratios of reduced and oxidized forms for individual redox couples, as is prominently exemplified for the redox pairs NAD^+^/NADH vs. NADP^+^/NADPH. Not least because of these aspects, it is unclear if at all, to which extent, and on which timescales chemical reduction of LOV receptors occurs inside of cells. This important aspect warrants further study in the future, e.g., by using fluorescent LOV derivatives^53,54^ with well-characterized flavin reduction midpoint potentials whose redox state can be tracked spectroscopically inside cells.

## Experimental section

### Molecular biology

For production of *A. sativa* phototropin 1 LOV 2 variants (*As*LOV2, residues 404-546, Uniprot O49003), the ampicillin resistance marker of an earlier pET-19b expression plasmid^16^ was replaced by a kanamycin (Kan) marker via Gibson cloning^55^. Within this vector, the *As*LOV2 variants are under control of a lactose-inducible T7 promoter and equipped with an N-terminal hexahistidine SUMO tag. To enhance expression levels, the T7 promoter and the translation initiation region governing *As*LOV2 expression were altered as recently described^56^. The resultant plasmid version is termed pET-19b+. *As*LOV2 variants were generated by site-directed mutagenesis according to the QuikChange protocol (Agilent Technologies). For expression of *N. crassa Vivid* (*Nc*VVD, residues 37-186, Uniprot Q9C3Y6), the pertinent gene was cloned into the above pET-19b vector conferring Kan resistance via Gibson cloning using an earlier construct as template^57^. *Nc*VVD variants were generated in this background by site-directed mutagenesis. The identity of all constructs was confirmed by Sanger DNA sequencing (MicroSynth, Göttingen, Germany).

### Protein expression and purification

*As*LOV2 variants were expressed and purified by adapting a previous protocol^16^. pET-19b or pET-19b+ expression vectors encoding a given *As*LOV2 variant were transformed into *Escherichia coli* CmpX13 cells^58^. Starter cultures containing 5 mL lysogeny broth (LB) medium, supplemented with 50 μg mL^-1^ kanamycin (LB/Kan), were inoculated with single bacterial clones and incubated overnight at 37°C. 4× 800 mL LB/Kan medium supplemented with 50 μM riboflavin were inoculated with the overnight starter cultures and incubated at 37°C and 225 rpm agitation in baffled flasks until an optical density at 600 nm of around 0.6 was reached. The temperature was lowered to 16°C, and 1 mM isopropyl β-D-1-thiogalactopyranoside were added. Incubation continued at 16°C and 225 rpm shaking for 3 days. Afterwards, cells were harvested by centrifugation and resuspended in buffer A (50 mM Tris(hydroxymethyl)amino-methane [Tris]/HCl pH 8.0, 150 mM NaCl, 20 mM imidazole) with protease inhibitor cocktail (complete ultra protease inhibitor kit, Merck, Darmstadt, Germany), 100 U DNAse I, and 0.1 g lysozyme added. Cells were lysed by sonication, and the supernatant after centrifugation was applied to a gravity-flow immobilized metal ion affinity chromatography (IMAC) column (HisPur cobalt resin, ThermoFisher). The column was washed with buffer B (50 mM Tris/HCl pH 8.0, 150 mM NaCl), followed by elution of bound protein with buffer C (50 mM Tris/HCl pH 8.0, 150 mM NaCl, 200 mM imidazole). Elution fractions were analyzed by denaturing polyacrylamide gel electrophoresis (PAGE) and pooled based on protein content and purity. The *As*LOV2 protein amount was gauged by absorbance spectroscopy, and 1 mg SUMO protease Senp2 was added per 150 mg *As*LOV2. After dialysis overnight into buffer B, the solution was passed again over an IMAC column, and the eluate was collected. To enhance purity, the eluate was again applied to the IMAC column. PAGE was used assess to sample purity. Certain *As*LOV2 variants required additional purification by anion-exchange chromatography on a Mono-Q resin. Finally, the sample was dialyzed into storage buffer D (20 mM Tris/HCl pH 8.0, 20 mM NaCl, 10% [w/w] glycerol), concentrated by spin filtration, aliquoted, and stored at –80°C.

The production of the *Nc*VVD variants proceeded similarly but with different buffers A (50 mM Tris/HCl pH 8.0, 150 mM NaCl, 10 mM imidazole, 10% [v/v] glycerol), B (50 mM Tris/HCl pH 8.0, 150 mM NaCl, 10% [v/v] glycerol), C (50 mM Tris/HCl Ph 8.0, 150 mM NaCl, 200 mM imidazole, 10% [v/v] glycerol), and D (20 mM Tris/HCl pH 8.0, 150 mM NaCl, 50% [v/v] glycerol).

### UV-vis absorbance spectroscopy

Spectra of dark-adapted *As*LOV2 and *Nc*VVD variants were acquired at 22°C on a diode-array absorbance spectrophotometer (Agilent Technologies, model 8453) equipped with Peltier temperature control (Agilent Technologies, model 89090A). The sample concentration was determined based on the absorbance maximum of the FMN cofactor around 450 nm using a molar extinction coefficient of 12,500 M^-1^ cm^-1^. For variants with incomplete chromophore loading, the concentration was determined based on the absorbance reading at 280 nm corrected for the contribution of the flavin chromophore. Spectra of the light-adapted LOV proteins were acquired after saturating exposure to a light-emitting diode (LED) with emission peaking at (470 ± 20) nm.

To record photoactivation kinetics, the LED was mounted in a custom 3D-printed adapter and set to an intensity of 3 mW cm^-2^. This approach ensures that all *As*LOV2 variants were assessed at the same illumination intensity and geometry. Dark-adapted samples were exposed to the LED light while recording absorbance spectra at periodic intervals. The decay of the absorbance signal at 450 nm, *A*_450_, with illumination time *t* was evaluated according to a single-exponential function (1):

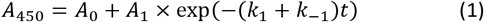

where *A*_0_ and *A*_1_ are amplitude terms, and *k*_1_ and *k*_-1_ denote the microscopic rate constants for photoactivation and dark recovery, respectively. Note that eq. (1) reflects that during photoactivation, the LOV variant already recovers to the resting state. The experimentally observable rate constant *k*_obs_ is thus the sum of the two microscopic rate constants. Data analyses and plotting were done with Fit-o-mat^59^. Once the light-adapted state was fully populated, the LED was switched off, and the recovery to the original, dark-adapted state was tracked by absorbance measurements and evaluated according to equation (2):

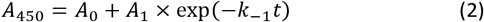

Experiments were repeated at several temperatures between 22°C and 38°C. The variation of *k*_-1_ with the absolute temperature *T* was analyzed according to the Arrhenius equation (3):

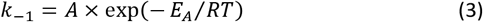

with *R* and *E*_a_ denoting the universal gas constant and the Arrhenius activation energy, respectively.

### Circular dichroism spectroscopy

The *As*LOV2 variants were analyzed by circular dichroism (CD) spectroscopy on a JASCO J-715 spectropolarimeter equipped with temperature control provided by a PTC-348WI Peltier element. CD measurements were conducted in 1-mm cuvettes at 22°C on samples containing 10 μM *As*LOV2 variant in 10 mM potassium phosphate pH 8.0, 10 mM NaCl. CD spectra between 190 nm and 250 nm were first recorded for the dark-adapted sample as an average of four consecutive scans. For the light-adapted state, the samples were exposed to blue light ([475 ± 20] nm, 50 mW cm^-2^) for 30 s immediately prior to each of four individual scans. The measured ellipticity was converted into molar ellipticity per residue, [*θ*]_MRW_, by taking into account the *As*LOV2 concentration.

The dark recovery of the *As*LOV2 variants after illumination ceases was followed by measuring [*θ*]_MRW_ at (208 ± 5) nm over time. The resultant data were evaluated according to eq. (2).

### Fluorescence spectroscopy

Measurements were performed at 22°C on an Agilent Cary Eclipse spectrometer equipped with Peltier temperature control. The *As*LOV2 variants were dissolved in buffer (20 mM Tris/HCl pH 8.0, 20 mM NaCl) to a concentration yielding an absorbance value at 450 nm of about 0.1 or less. Emission spectra were recorded between 450 and 700 nm at an excitation wavelength of (435 ± 10) nm and using a bandwidth of 5 nm for emission. Excitation spectra were recorded between 350 and 530 nm at 5 nm bandwidth for an emission wavelength of (550 ± 10) nm.

The fluorescence quantum yield, *Φ*_F_, of the *As*LOV2 variants was determined based on comparison to the reference fluorophore coumarin 153 with known *Φ*_F_ in ethanol of 54% ^60^. For the measurements, dilutions of the *As*LOV2 variant and the reference dye, respectively, were prepared to yield absorbance values of 0.02, 0.04, 0.06, 0.08, and 0.1 at the excitation wavelength of 435 nm. Fluorescence emission spectra were recorded between 450 and 700 nm and integrated. The integrated fluorescence intensities were plotted against the absorbance of the solution and evaluated by linear fitting to determine the slopes *m*_F_ and *m*_r_ for a given *As*LOV2 variant and the reference dye, respectively (see Fig. 3d). The fluorescence quantum yield *Φ*_F_ of a given *As*LOV2 variant can then be calculated according to equation (4):

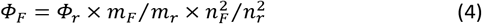

where *Φ*_r_ denotes the fluorescence quantum yield of the reference, and *n*_F_ and *n*_r_ refer to the refractive indices of the buffer and ethanol used for the *As*LOV2 variant and the reference fluorophore, respectively.

### Measurements of reduction midpoint potential

To determine the reduction midpoint potential *E*_0_ for the two-electron reduction of the flavin quinone to hydroquinone, we dissolved between 20 and 40 μM of a given *As*LOV2 variant in buffer (50 mM sodium phosphate pH 7.0). To facilitate electron exchange between solution and analyte, methyl viologen was added at a concentration of 10 μM. To allow determination of *E*_0_, the redox indicator phenosafranine with a known midpoint potential *E*_r0_ of –252 mV was included in the solution at 6 μM concentration. Oxygen was removed by adding an oxygen scavenger system comprising 10 mM glucose, 50 μg mL^-1^ glucose oxidase, and 5 μg mL^-1^ catalase. In addition, prior to the measurement, the solution was flushed with nitrogen under constant stirring. The reduction was initiated by adding 250 mM dithiothreitol (buffered at pH 7.0), immediately followed by layering paraffin oil over the aqueous solution to hinder oxygen re-entry. The reaction was followed by recording absorbance spectra on a SEC 2020 diode-array spectrophotometer (ALS, Tokyo, Japan) over several hours until complete reduction occurred. Mock experiments leaving out DTT confirmed that the probe light used for the measurement does not photoactivate the *As*LOV2 nor the *Nc*VVD variants to significant extent.

From the absorbance data at each timepoint, the fractions *f*_red_ and *f*_ox_ of the given *As*LOV2 variant in its reduced and oxidized states were calculated based on reference spectra of fully reduced and fully oxidized *As*LOV2 variant determined in control experiments. Likewise, for each timepoint, the corresponding fractions *f*_r,red_ and *f*_r,ox_ of the reference dye were calculated. According to the Nernst equation (5), the fractions of reduced and oxidized species depend on the reduction potential *E* and their reduction midpoint potential *E*_0_ as follows:

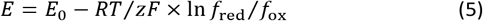

where *F* is the Faraday constant and *z* the number of transferred electrons, i.e., *z* = 2 for both the reduction of the *As*LOV2 variant and the phenosafranine reference. Provided the reduction reaction proceeds sufficiently slowly to ensure constant redox equilibration throughout the system, the current reduction potential *E* at a given time can hence be inferred from the reference dye. By equating the Nernst equations for the *As*LOV2 variant and the reference dye, one obtains equation (6):

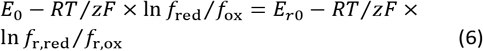

As *z* is equal for phenosafranine and flavin reduction, this equation simplifies to equation (7):

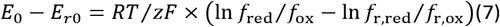

The difference in the midpoint potentials of analyte and dye can thus be determined as the axis offset of the linear correlation between the terms for analyte and dye, respectively, on the righthand side of eq. (7). As is apparent from eq. (7), the slope of the correlation should be unity, and deviations form this value may hint at insufficient redox equilibration. The slope generally ranged between 0.8 and 1.2, and experiments outside this interval were disregarded. The evaluation was performed using Fit-o-mat^59^. Redox titrations were repeated at least two times, and the *E*_0_ values for each variant were reproducible within ± 3 mV.

### Sequence analyses

To gauge the distribution of methionine residues in natural LOV receptors, we performed sequence searches similar to before^16,18^. In brief, a BLAST search against GenBank with *As*LOV2 (residues 404-519) as the query sequence yielded 76,228 hits at an *E*-value of cutoff 10. We filtered these for entries possessing at minimum eight of nine residues conserved in LOV receptors^61^ (corresponding to residues G447, N449, C450, R451, F452, L453, Q454, N482, N492, and Q513 in *As*LOV2), followed by clustering at a 90% identity level using USEARCH^62^. Next, the resulting 10,017 centroid sequences were aligned with MUSCLE^63^ and analyzed for the relative frequency of methionine residues at individual positions within the resulting multiple sequence alignment. These frequencies were converted into a color code and plotted onto the *As*LOV2 structure (pdb entry 7pgx)^16^.

## Author Contributions

**A.G.F**. – Conceptualization, Formal Analysis, Investigation, Methodology, Visualization, Writing – original draft. **A.M**. – Conceptualization, Formal Analysis, Funding acquisition, Supervision, Visualization, Writing – original draft, Writing – review & editing.

## Acknowledgements

We thank U. Krauß, T. Krebel, and L. Ott for comments on the manuscript, our laboratory colleagues for discussion and the Deutsche Forschungsgemeinschaft for funding (MO2192/8-1 to A.M.).

## Supplementary Material

**Supplementary Figure 1.**
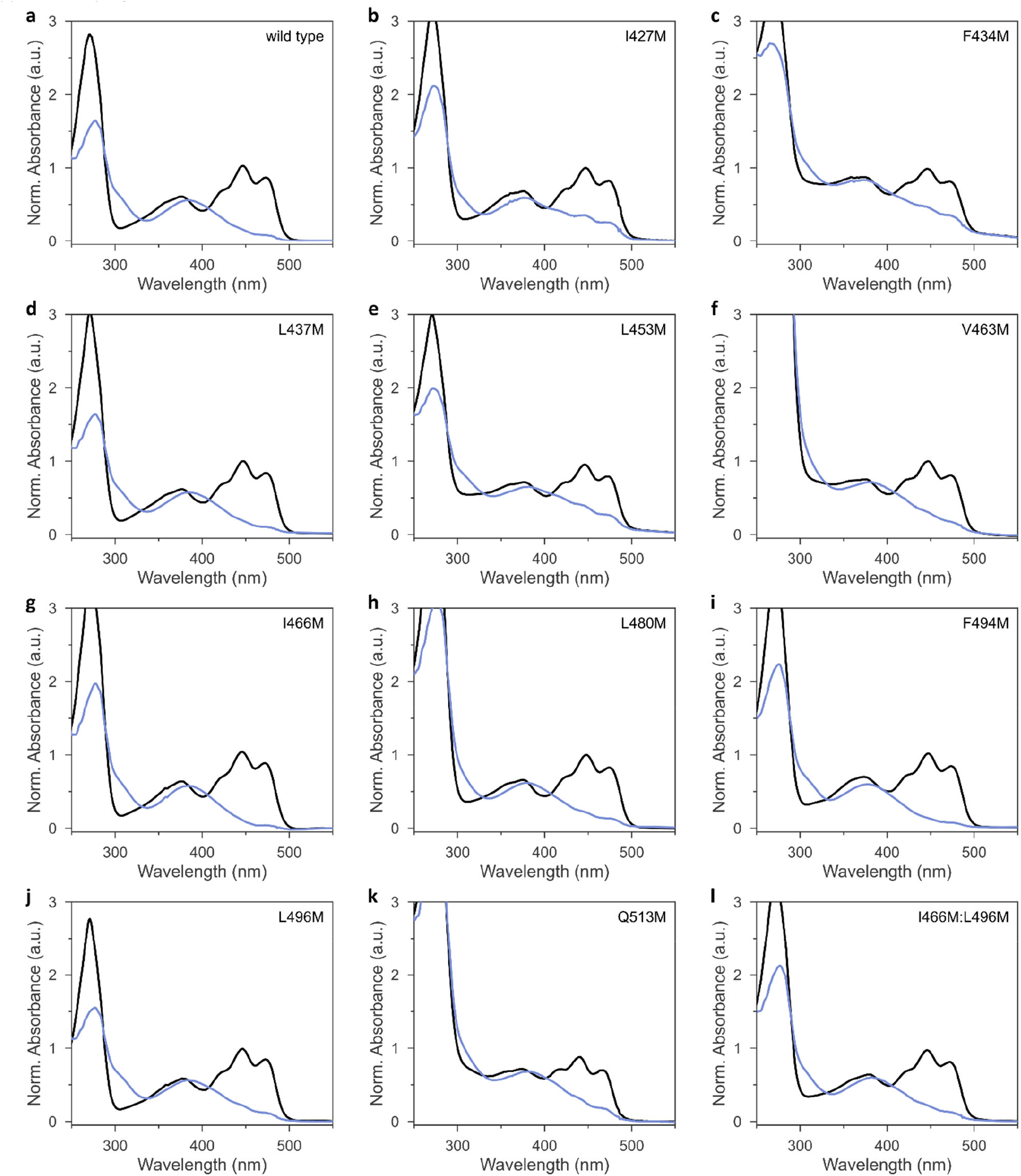
UV-vis absorbance spectra of the *As*LOV2 variants in their dark-adapted (black lines) and light-adapted states (blue). **a**, *As*LOV2 wild type. **b**, *As*LOV2 I427M. **c**, *As*LOV2 F434M. **d**, *As*LOV2 L437M. **e**, *As*LOV2 L453M. **f**, *As*LOV2 V463M. **g**, *As*LOV2 I466M. **h**, *As*LOV2 L480M. **i**, *As*LOV2 F494M. **j**, *As*LOV2 L496M. **k**, *As*LOV2 Q513M. **l**, *As*LOV2 I466M:L496M.

**Supplementary Figure 2.**
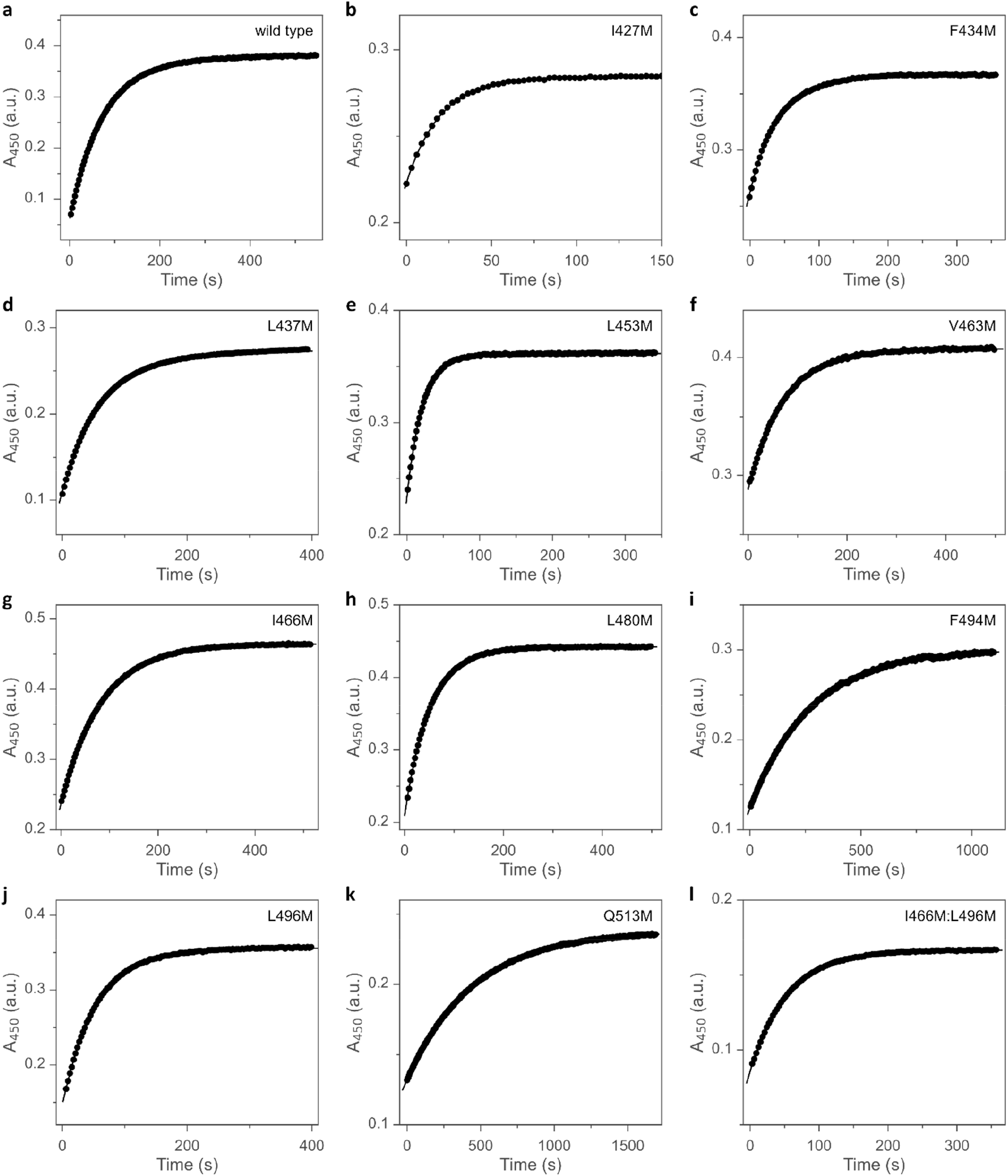
Dark-recovery kinetics of the *As*LOV2 variants monitored by absorbance measurements at 450 nm. **a**, *As*LOV2 wild type. **b**, *As*LOV2 I427M. **c**, *As*LOV2 F434M. **d**, *As*LOV2 L437M. **e**, *As*LOV2 L453M. **f**, *As*LOV2 V463M. **g**, *As*LOV2 I466M. **h**, *As*LOV2 L480M. **i**, *As*LOV2 F494M. **j**, *As*LOV2 L496M. **k**, *As*LOV2 Q513M. **l**, *As*LOV2 I466M:L496M.

**Supplementary Figure 3.**
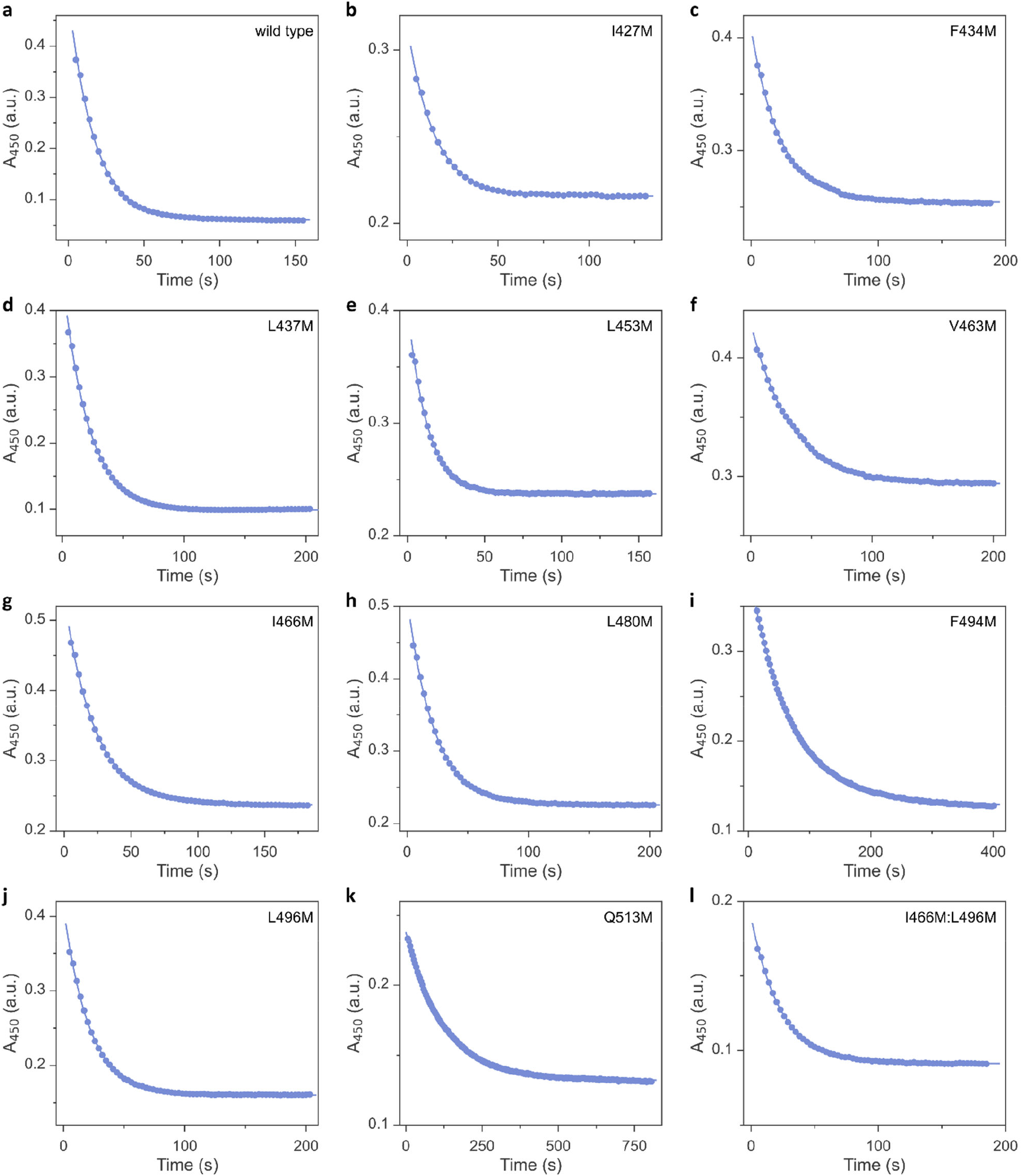
Photoactivation kinetics of the *As*LOV2 variants monitored by absorbance measurements at 450 nm. A set illumination geometry and intensity (3 mW cm^-2^) was used in all measurements. **a**, *As*LOV2 wild type. **b**, *As*LOV2 I427M. **c**, *As*LOV2 F434M. **d**, *As*LOV2 L437M. **e**, *As*LOV2 L453M. **f**, *As*LOV2 V463M. **g**, *As*LOV2 I466M. **h**, *As*LOV2 L480M. **i**, *As*LOV2 F494M. **j**, *As*LOV2 L496M. **k**, *As*LOV2 Q513M. **l**, *As*LOV2 I466M:L496M.

**Supplementary Figure 4.**
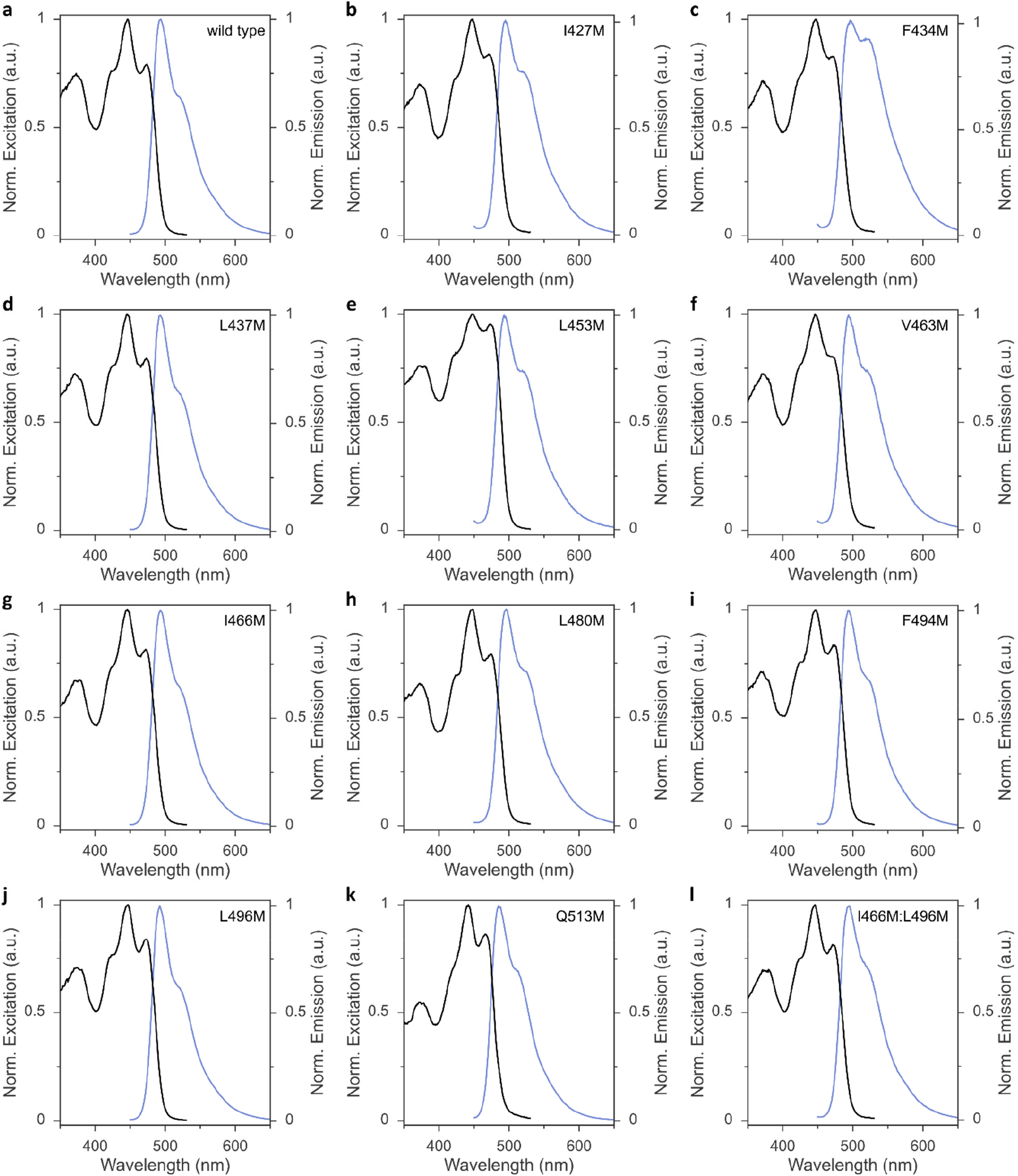
Fluorescence excitation (black lines) and emission spectra (blue) of the *As*LOV2 variants. **a**, *As*LOV2 wild type. **b**, *As*LOV2 I427M. **c**, *As*LOV2 F434M. **d**, *As*LOV2 L437M. **e**, *As*LOV2 L453M. **f**, *As*LOV2 V463M. **g**, *As*LOV2 I466M. **h**, *As*LOV2 L480M. **i**, *As*LOV2 F494M. **j**, *As*LOV2 L496M. **k**, *As*LOV2 Q513M. **l**, *As*LOV2 I466M:L496M.

**Supplementary Figure 5.**
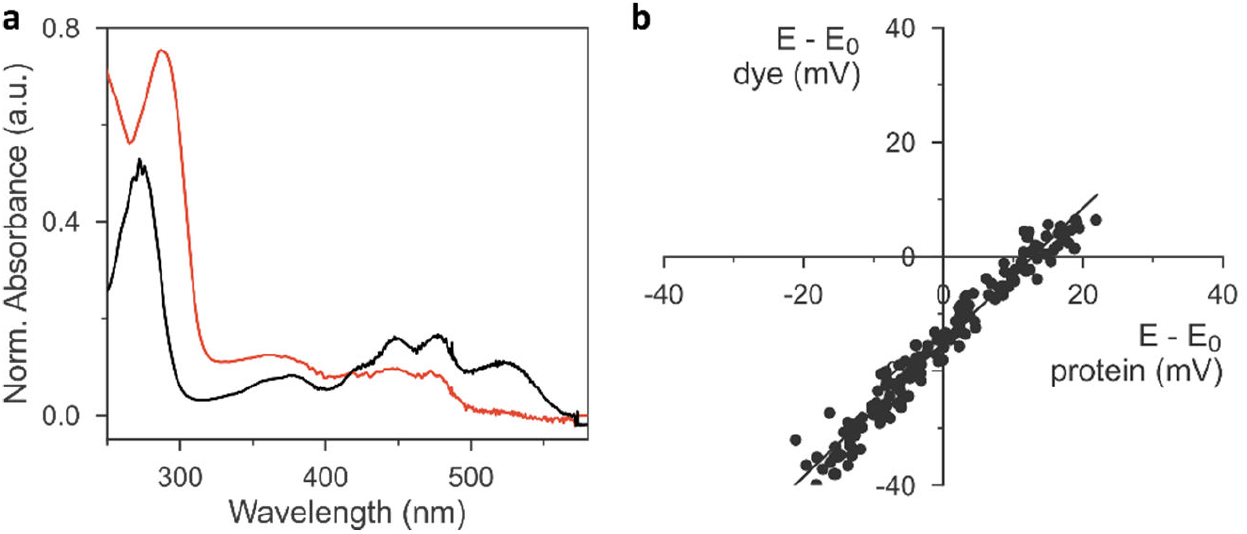
Determination of the flavin reduction midpoint potentials in *As*LOV2 wild type by the xanthine-oxidase (XO) method. **a**, Absorbance spectra at the start of the XO reduction kinetics (black lines) and after 5 hours (red). **b**, The correlation of the oxidized and reduced fractions of the redox indicator phenosafranine and *As*LOV2 according to the Nernst equation allows the determination of the flavin reduction midpoint potential in *As*LOV2. The *y* axis denotes *E* – *E*_0,r_ for the dye, and the *x* axis plots the corresponding values *E* – *E*_0_ for the LOV protein.

**Supplementary Figure 6.**
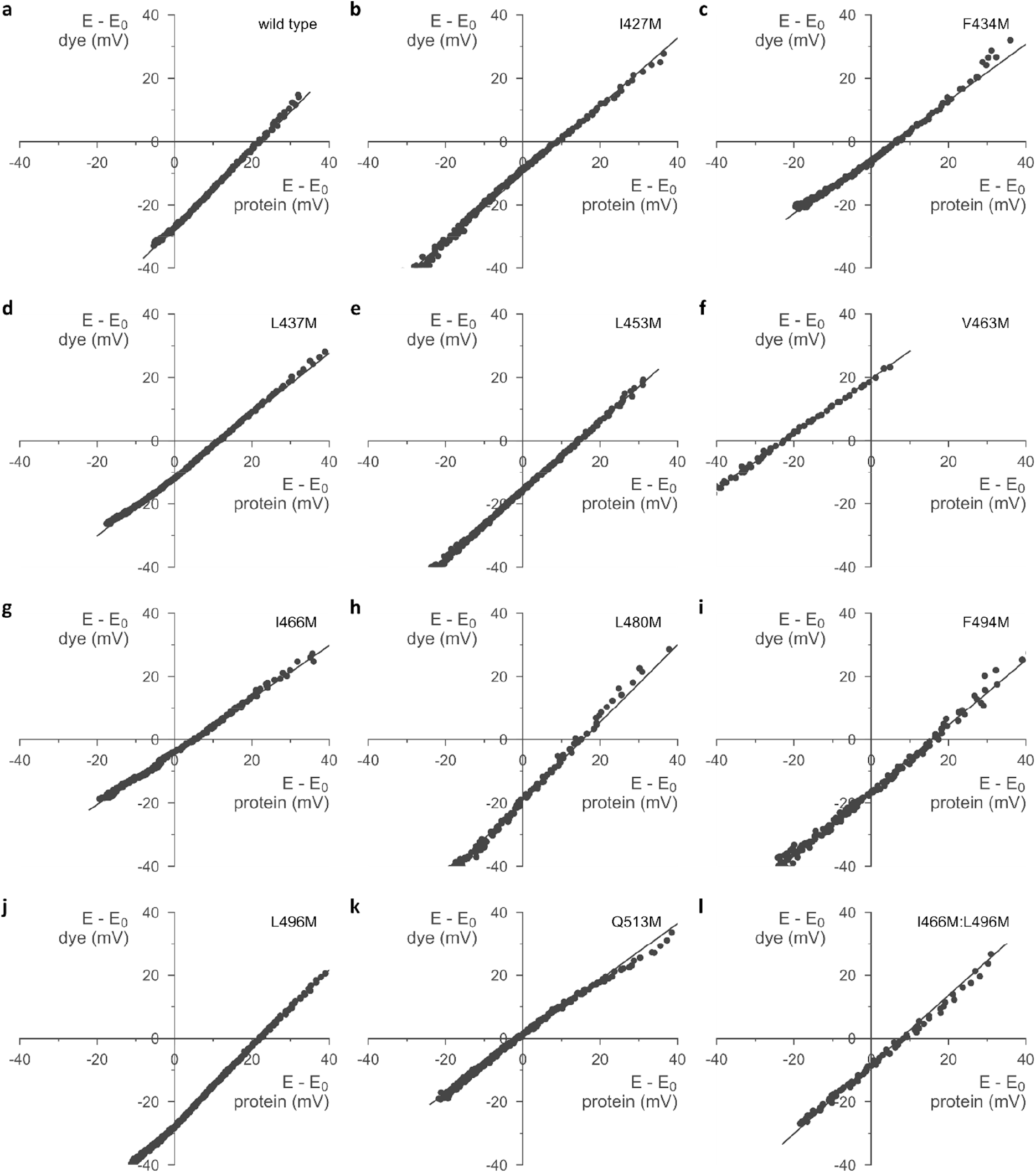
Determination of the flavin reduction midpoint potentials in the *As*LOV2 variants. The *y* axis denotes *E* – *E*_0,r_ for the dye, and the *x* axis plots the corresponding values *E* – *E*_0_ for the LOV protein. **a**, *As*LOV2 wild type. **b**, *As*LOV2 I427M. **c**, *As*LOV2 F434M. **d**, *As*LOV2 L437M. **e**, *As*LOV2 L453M. **f**, *As*LOV2 V463M. **g**, *As*LOV2 I466M. **h**, *As*LOV2 L480M. **i**, *As*LOV2 F494M. **j**, *As*LOV2 L496M. **k**, *As*LOV2 Q513M. **l**, *As*LOV2 I466M:L496M.

**Supplementary Figure 7.**
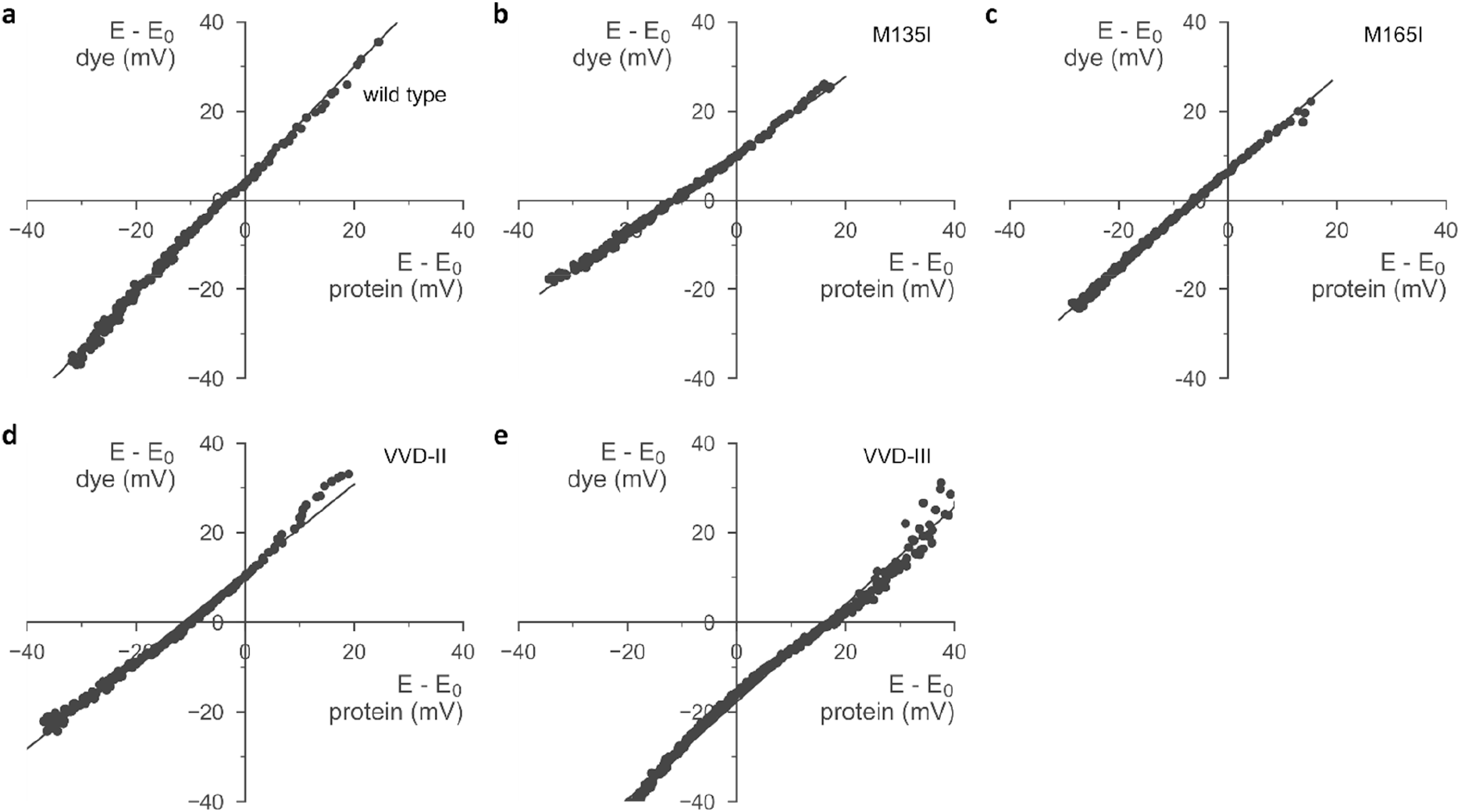
Determination of the flavin reduction midpoint potentials in the indicated *Nc*VVD variants. The *y* axis denotes *E* – *E*_0,r_ for the dye, and the *x* axis plots the corresponding values *E* – *E*_0_ for the LOV protein. **a**, *Nc*VVD wild type. **b**, *Nc*VVD M135I. **c**, *Nc*VVD M165I. **d**, *Nc*VVD-II (M135I:M165I). **e**, *Nc*VVD-III (C108A:M135I:M165I).

**Supplementary Figure 8.**
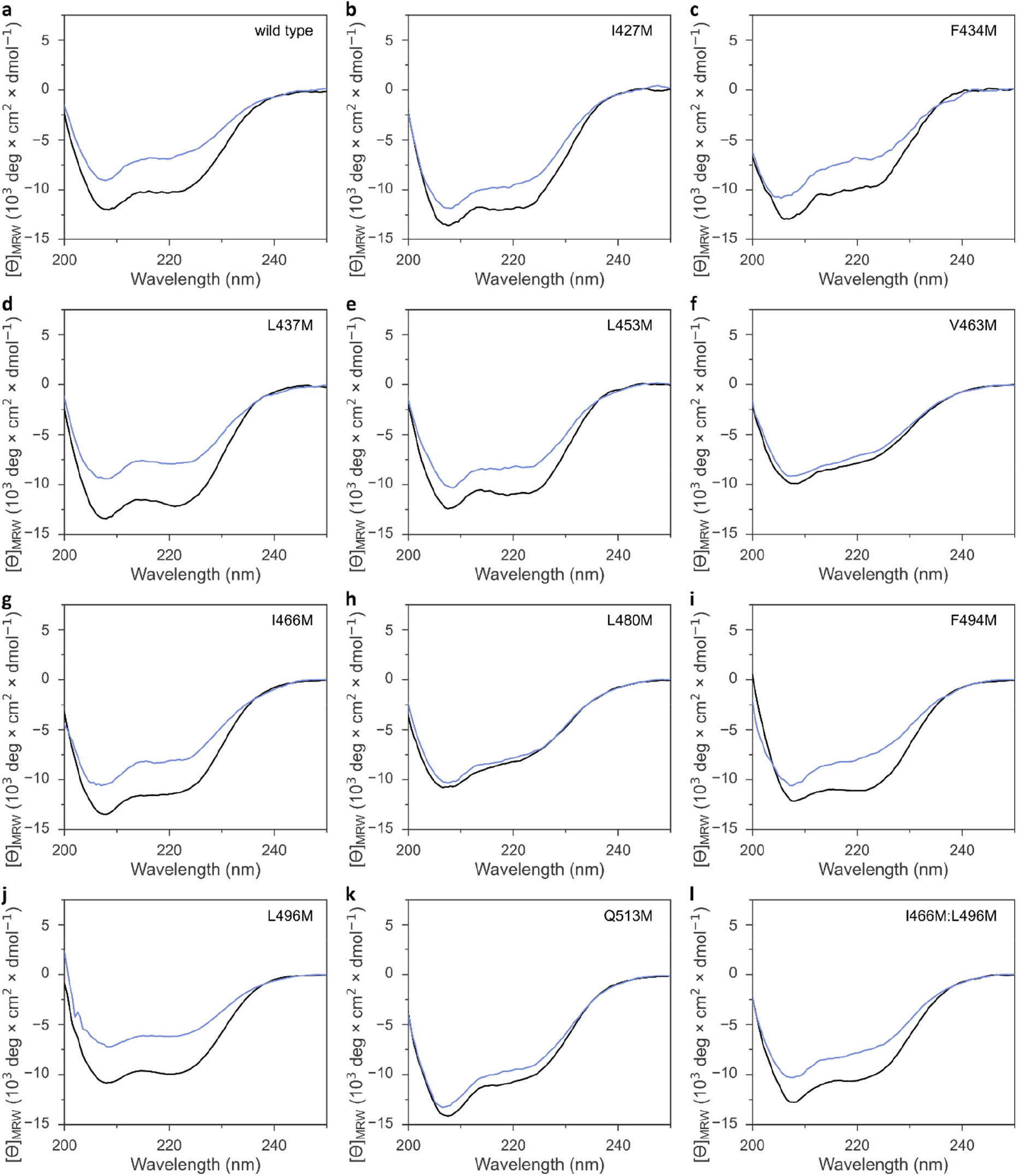
Far-UV circular dichroism spectra of the *As*LOV2 variants in their dark-adapted (black lines) and light-adapted states (blue). **a**, *As*LOV2 wild type. **b**, *As*LOV2 I427M. **c**, *As*LOV2 F434M. **d**, *As*LOV2 L437M. **e**, *As*LOV2 L453M. **f**, *As*LOV2 V463M. **g**, *As*LOV2 I466M. **h**, *As*LOV2 L480M. **i**, *As*LOV2 F494M. **j**, *As*LOV2 L496M. **k**, *As*LOV2 Q513M. **l**, *As*LOV2 I466M:L496M.

